# qPyCR: A qPCR Data Analyzer for Template Quantification and the Detection of Performance Outliers

**DOI:** 10.64898/2026.10.08.757329

**Authors:** Sean D. Moore

**Affiliations:** Burnett School of Biomedical Sciences, University of Central Florida, Orlando, FL 3286, USA; Genomics and Bioinformatics Cluster, University of Central Florida, Orlando, FL 3286, USA

## Abstract

**Motivation:** Real-time qPCR data are commonly analyzed to obtain a Cq values, which are used to compare the relative abundance of a targeted template. However, improper background subtraction distorts Cq and changing the assigned threshold can alter experimental conclusions. In addition, a Cq values provide no information on the quality of the qPCR for a given sample. Alternatively, global fitting of qPCR data provides agnostic quantification without the use of Cq or thresholds, and the fitting terms provide a useful metric of reaction performance. Current implementations of global fitting are limited to custom in-house programs.

**Results:** PyCR is comprehensive analysis tool written in Python that performs background signal subtractions, computes a threshold, and reports Cq values in a traditional manner. The program also performs global fitting and reports template abundances along with reaction quality values. The quality values can be used to assign performance standards, which allow researchers to distinguish aberrant reactions from true changes in template abundance.

**Availability and Implementation:** Code and example data are freely available on GitHub (https://github.com/sdmoore-labs/qPyCR) and on Zenodo (https://doi.org/10.5281/zenodo.21115342).

**Supplementary information:** Supplementary data are available at *bioRxiv* online.

## Introduction

Quantitative, real-time polymerase chain reactions (qPCR) are widely used to establish the relative abundance of nucleic acid templates. The accumulation of PCR amplicons after each thermal cycle is monitored by fluorescence signals, which increase as new DNA accumulates and the signals eventually plateau as each reaction subsides (Jansson and Hedman, 2019). The thermal cycles at which the accumulation of amplicons becomes detectable is proportional to the amount of starting template, so the quantification of templates is inferred from the thermal cycle at which an amplification signal reaches a set threshold value (referred to as Ct or Cq) (Bustin et al., 2025; Higuchi et al., 1993; Pfaffl, 2001).

For diagnostic procedures, it is common practice to use a predetermined cycle number as the boundary between template-positive (lower Cq) and template-negative (higher Cq) samples. In practice, operational differences between experiments, software, or equipment will cause variations in Cq calling, which can shift the positive/negative sample calls (Bustin et al., 2025; Ruiz-Villalba et al., 2021). Moreover, the threshold positions used to calculate Cq are not standardized, in part because different qPCR methods and platforms generate arbitrary fluorescence values and also because the algorithms used to assign thresholds may be different or proprietary. On top of platform variation is the concerning capability of end-users to manually adjust a threshold; so, it is possible to shift a collection of samples from positive to negative (or *visa versa*) (Ruiz-Villalba et al., 2021). Despite continued efforts by the community to encourage best practices and the submission of raw qPCR data during review and publication (Bustin, 2024; Hampton et al., 2025; Ruijter et al., 2015; Untergasser et al., 2023), many reports still only include Cq values and selected threshold values are rarely reported.

### Background subtraction

The fluorescent signals arising from amplicon accumulation are recorded along with a substantial background fluorescence, which varies among samples and requires subtraction before subsequent analysis. Inaccurate background subtraction significantly impacts Cq calculations (Ramakers et al., 2003; Ruijter et al., 2009; Ruiz-Villalba et al., 2021), and background-subtraction algorithms implemented by commercial software can dramatically distort data sets by additionally tilting (rotating) or artificially flattening the data in an attempt to create a near-zero baseline in early cycles. Although several third-party tools are available that can correct for backgrounds using different approaches (Pabinger et al., 2014), there is no standard implementation across platforms. The rationale for such adjustments is the contention that qPCR data should remain near zero until amplification is observed; however, detectable amplification can be present in early-cycle data and the applied background adjustments can create unrealistic datasets.

### Cq calling

After background subtraction, qPCR data are usually log-transformed to reveal the exponential amplification regions (log-linear). Those regions are then fit using linear regression and a single threshold value (fluorescence intensity) is selected that crosses each sample’s log-linear region. Cq is assigned as the fractional cycle number at which the log-linear fits cross the threshold value (Higuchi et al., 1993). This approach has been implemented since the earliest use of qPCR and it remains as a stapled convention in data analysis (Bustin et al., 2025; Ruijter and van den Hoff, 2025; Untergasser et al., 2023).

### Reaction outliers

One drawback of qPCR is measurement variation, which commonly involves Cq values from replicate measurements of the same sample being scattered by ∼0.5 cycles or more (corresponding to over a ∼1.5-fold deviation in apparent quantity). To address these variations, Cq values from replicate measurements are averaged before subsequent analysis (Ruiz-Villalba et al., 2021). Measurement distortions that create reaction outliers can arise from mis-loading, unseated wells, sample evaporation, or optical interference (so-called a ‘bad’ wells). Such outliers among a set of technical replicates can be identified as having deviations in the overall reaction profile relative to the set (Rödiger et al., 2015), but there is currently no way to determine if an entire replicate set is deviant. Moreover, when Cq values are used to establish target concentrations relative a standard reference, template-concentration-dependent changes in PCR efficiency obscure the relationship between ΔCq and target abundance (H.Hampton et al., 2025). There is no current method that can distinguish altered reaction performance from *bona fide* changes in template abundance.

### Global fitting

Global fitting of qPCR data involves fitting all of a reaction’s qPCR data points to a PCR model. In prior work, we developed a PCR model that accurately describes experimental qPCR data and we demonstrated that the fitted models can be used to compute relative template abundance (Carr and Moore, 2012). Because this global fitting approach does not use Cq or a threshold, small errors in background subtraction are less impactful and the abundance calculations are not influenced by user settings. Recently, I reported that the ratio of the two variables obtained by our global fitting model appears to reflect the performance quality of a given reaction (Moore, 2025). This discovery motivated the development pPyCR so that global fitting and quality control will be more accessible. qPyCR accepts raw (unadjusted) qPCR data, subtracts backgrounds, computes Cq, and applies global fitting to establish template abundance and to evaluate reaction performance.

## Methods

### Experimental conditions

Commercial qPCR reagent mixtures were used to generate the data analyzed in each figure. Details of each reaction condition are described in ReadMe files located in their respective data folders (**Supplementary Data SD1**). Oligonucleotide primer sequences and descriptions are provided in **Supp. Data SD2**.

### Software and data analysis

Fluorescence data were converted to CSV or TSV format using Excel (Microsoft). Per-cycle PCR efficiency calculations for Cq *vs*. Log_template plots were derived from linear regression slopes (*m*) using Excel: *E* = ((10^(-1/*m*)) - 1) (Bustin et al., 2025). Data plotting and statistical analyses were performed using Prism 10 (Graphpad). Figures were generated using Illustrator (Adobe Systems). The qPyCR program was constructed in Python 3.+ using Google Colaboratory (https://colab.research.google.com). Python code cells were written using the Cursor IDE with the assistance of its ‘agent mode’ (https://cursor.com/).

### Data input

qPyCR accepts either pre-adjusted or unadjusted ‘raw’ data from a qPCR platform in a simple CSV format, with the first column containing the cycle numbers and subsequent columns containing the sample/well IDs and the fluorescence values. Alternative input formats include Real-time PCR Data Essential Spreadsheets in TSV format (RDES) and Real-time PCR Data Markup Language file (RDML) (Ruijter et al., 2015; Untergasser et al., 2023). The method for viewing and exporting raw data from some platforms may be inconspicuous. For example, on the Bio-Rad platforms used in this study, raw data were obtained by: switching to the ‘Quantification Data’ tab, selecting ‘RFU’ in the dropdown menu, selecting ‘Settings’ from the main menu bar, and changing ‘Baseline Setting’ to ‘No Baseline Subtraction’. The data were then copied and pasted into a spreadsheet.

### Data outputs

The requested outputs are assigned during the data loading step: default = a single output file that contains sample IDs, Cq, and global fitting values ([input]_qPyCR_Analysis_Outputs_[date]_[time].csv). The evaluation mode (-e) = yields the default output in addition to files containing the background-adjusted data (final_adjusted_data_*.csv), a plot of the Cq/threshold assignments (log_with_threshold_*.png), and a plot of the global fitting (Global_fitting_plot_*.png). The debug mode (-d) provides the prior files in addition to numerous data and plot files from each stage of the pipeline for each sample.

## Results

### qPyCR overview

A summary of the operations performed by qPyCR are shown in Figure 1. The program accepts either raw or platform-adjusted qPCR data, but unadjusted data are recommended. After adjusting the data to remove background (default: True), a flag is applied to each reaction indicating whether or not there was substantial amplification based on the signal-to-noise of the final fluorescence compared to the baseline. The data are then globally fit using the PCR model to obtain *max* and *K*^*D*^ for each reaction. An optional baseline refinement stage follows (default: False) that attempts to fine-tune the background adjustment by iteratively shifting the subtracted background value and re-fitting the data to the PCR model to minimize fitting residuals. In pilot testing, it was found that this approach tended to shift some baselines away from zero, so it is recommended to only use this stage to interrogate data for its overall conformance to an ideal PCR model. After background subtraction, the data are processed to identify linear regions in their exponential phases, a threshold is assigned, and Cq values are calculated. The threshold is assigned as the median fluorescence value of all exponential windows in the analyzed set. The next stages use fixed *max* and *K*^*D*^ values to model an entire reaction from a cycle 0 *seed* value. Because *seed* values of an experiment can cover a very large range, an initial guess of each *seed* is first calculated using the Cq. The *seed* value is then optimized by iteratively adjusted it to minimize the fitting residuals. The Cq, optimized *seed, max, K*^*D*^, and *max*/*K*_*D*_ are exported as the default output.

**Figure 1.**
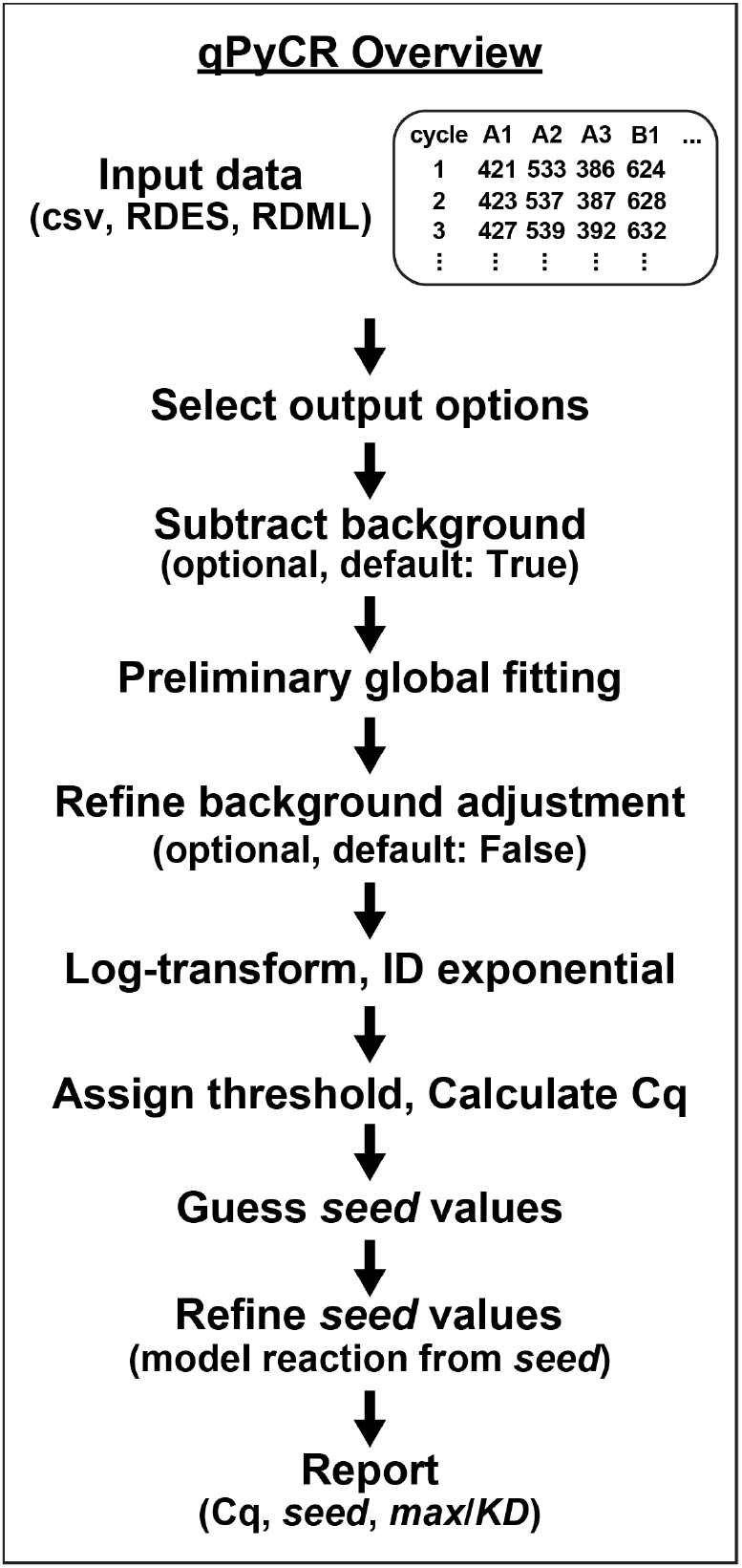
Overview of qPyCR operations. A flowchart highlighting the data processing operations performed by qPyCR.

### Adjusting data to remove background

qPyCR applies a dynamic background subtraction algorithm that can process data from early-amplifying samples that do not exhibit a linear baseline. It initially attempts fitting early cycle data to a simple exponential function, with constraints on signal variance and the fitted exponent to avoid noisy or unrealistic amplification data. If exponential fitting fails, it reverts to a dynamic linear regression through the early data. The program attempts to extend the number of included data points for longer regression windows through later cycles while monitoring for trend deviations that may indicate entry into the amplification phase, at which point the expansion of the regression window is halted. After exponential or linear regression fitting, the data are adjusted by subtracting the computed cycle 0 fluorescence values and the linear regression slopes are used to level those baselines.

The qPyCR background adjustments outperform a common commercial platform when dealing with data having long baselines (late amplification, **Fig. 2A-D**) and on data having no baseline (early amplification, **Fig. 2E-H**). In each case, the qPyCR-adjusted data trend toward zero fluorescence at cycle 0, which is a realistic expectation after background subtraction.

**Figure 2.**
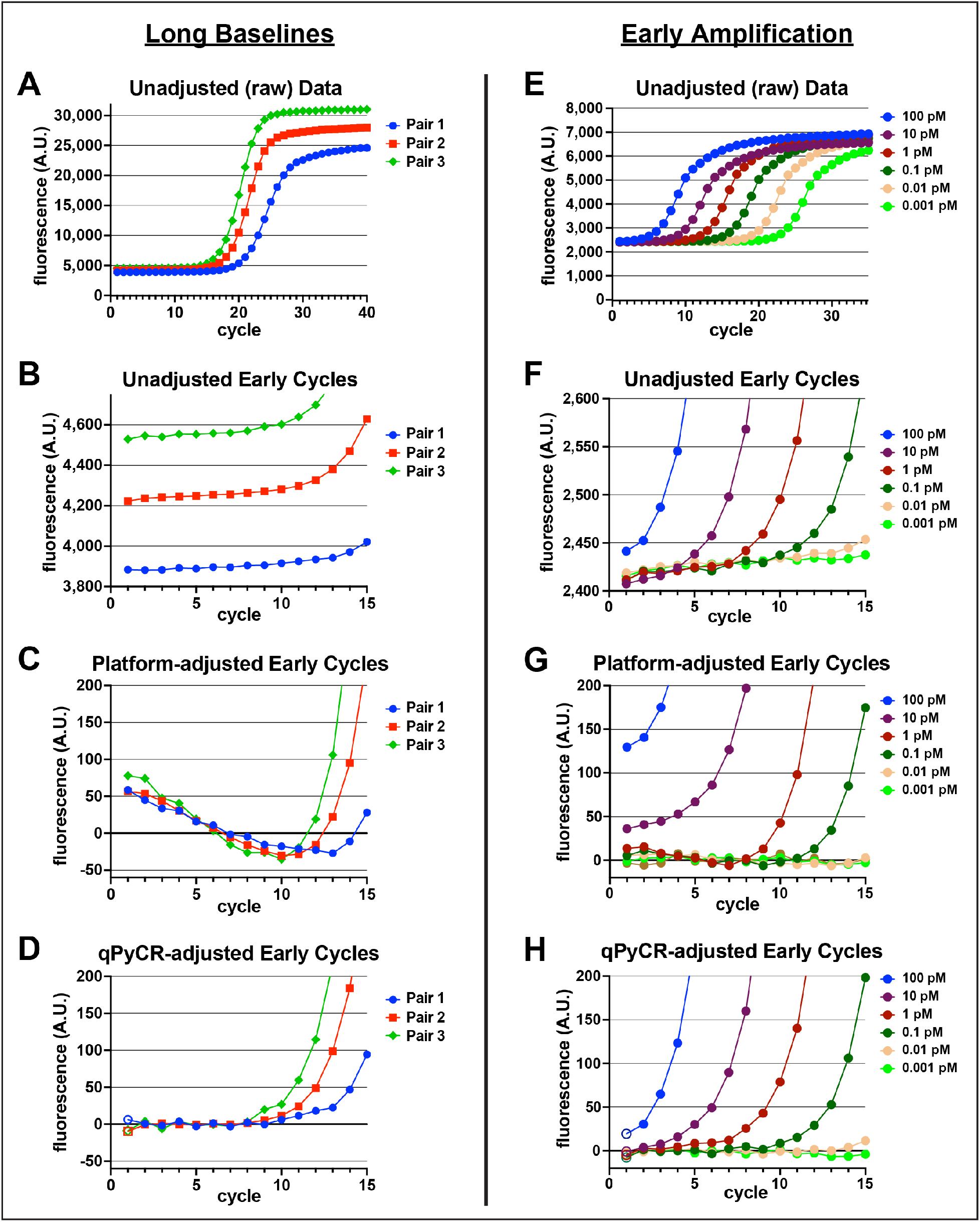
Background subtraction examples. Two sets of qPCR data were processed using a commercial platform (Bio-Rad) or by qPyCR. The *left panels* contain data from an experiment comparing the performance of three different primer pairs (*Pair 1-3*) under the same thermal-bias PCR conditions targeting a section of the *E. coli* 16S rRNA gene using 5 μg/mL genomic DNA. **A)** Examples of the raw data are plotted (one curve displayed from three replicates each for clarity). **B)** A plot of the raw data from early cycles showing that these reactions exhibited long baselines. **C)** A plot of data from the early cycles after platform adjustment reveals unrealistic downward trends of the baselines. **D)** Early cycle data after adjustment by qPyCR have level baselines that trend toward zero at cycle 0. The *right panels* contain data from a standard dilution series generated using a commercial quantification kit (NEBNext). **E)** Examples of the raw data (one curve from three replicates each). **F)** Inspection of the unadjusted early cycles reveals early-amplifying samples lacking baselines. **G)** The platform-adjusted data do not trend toward zero and the relative spacing of the curves is distorted. **H)** The qPyCR-adjusted data trend toward zero and the curve spacing reflects that of the raw data. The cycle 1 data were plotted using open symbols because this version of qPyCR skips the first data point while establishing background.

### Calculating relative abundance and evaluating reaction performance

The datasets from Fig. 2 were processed by the qPCR platform to determine Cq values, and also processed by qPyCR to determine Cq, abundance (*seed*), and performance (*max*/*K*_*D*_). The Cq values calculated by qPyCR mirror those generated by the platform, but they were shifted by ∼5 cycles for the primer evaluation data (**Fig. 3A**) and by ∼2.5 cycles for the commercial standards (**Fig. 3D**). These substantial shifts highlight the impact of differences in background subtraction and threshold assignment procedures (Ruiz-Villalba et al., 2021).

**Figure 3.**
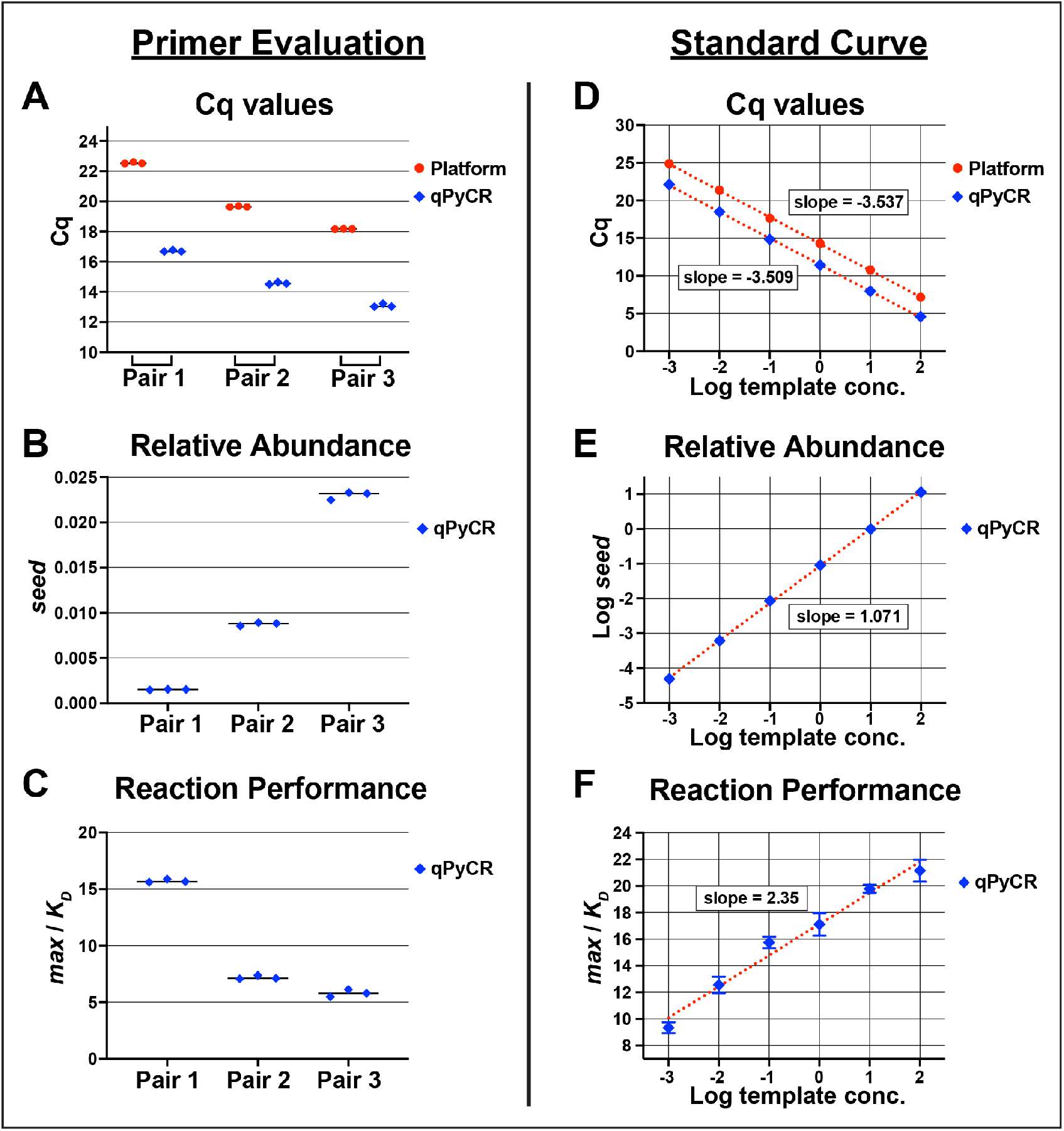
Quantification and reaction performance evaluations. The two experimental qPCR data sets shown in Figure 1 were evaluated for quantification (Cq and *seed*) and performance (*max*/*K*_*D*_). The left panels compare data generated using primer pairs 1-3 under the same conditions. **A)** Cq values reported either by the platform (Bio-Rad) or by qPyCR. **B)** Apparent relative target abundance was established using the *seed* values (arbitrary units). **C)** Relative reaction performance was compared using *max*/*K*_*D*_ values. The right panels show the analysis of the NEBNext standard curve reactions. **D)** Plots of Cq *vs*. Log(template concentration) were fit using linear regression to obtain slopes (platform data: slope = -3.537, R^2^ = 0.9995; qPyCR data: -3.509, R^2^ = 0.9994), which correspond to calculated per-cycle reaction efficiencies of 91.8% and 92.8%, respectively. **E)** Linear regression of *seed vs*. Log(template concentration) indicates a strong correlation (slope = 1.071, R^2^ = 0.9997). **F)** Template concentration-dependent reaction performance of the standards was evaluated as *max*/*K*_*D*_ *vs*. Log(template concentration). The performance of these standards exhibited a positive correlation with template abundance (slope = 2.35, R^2^ = 0.979, residual standard deviation (Sy.x) = 0.719); these values can be used to set performance expectations for experimental measurements falling withing the concentration range of the standards.

Global fitting to our PCR model returns the values of two variables (*max* and *K*^*D*^) that describe the shape of a qPCR amplification curve (Carr and Moore, 2012). Once obtained, a modeled amplification curve using those numbers is used to establish a ‘*seed*’ value, which represents the fluorescence signal at cycle 0 (the signal from the target template before amplification). Unlike Cq, *seed* values are linearly related to template abundance. The *seed* values computed by qPyCR for the primer evaluation tests revealed that the reactions generated using Pair 3 apparently had more template (**Fig. 3B**); however, each reaction in that comparison used the same amount of template, so that difference reflects a more efficient amplification. The *seed* values obtained from qPyCR analysis of the commercial standards had a nearly 1-to-1 correlation with the input template amounts (**Fig. 3E**), which confirms *seed* values can be used to establish relative template abundance with high accuracy.

The ratio of *max*/*K*_*D*_ was discovered to reflect qPCR reaction performance, with lower ratios suggesting more efficient reactions (Moore, 2025). Consistent with this idea, when the primer evaluation data were compared using this metric, *max*/*K*_*D*_ for Pair 2 and Pair 3 were similar and notably lower than that of Pair 1 (**Fig. 3C**). The *max*/*K*_*D*_ analysis of the commercial standards data revealed a sloped linear trend, with higher template concentrations yielding higher *max*/*K*_*D*_ values (**Fig. 3F**). This finding indicates that the reaction performance of the PCR standards was not consistent, despite the fact that the Cq and *seed* values were well-correlated with template concentration. Trends and scatter in the performance values of standards can be used to establish confidence intervals for experimental reaction performance. In this example, experimental performance values deviating by more than ∼2.5 units from the standards near that template concentration may indicate an issue with reaction quality. It is important to note that because *max*/*K*_*D*_ ratios are derived from arbitrary fluorescence values from a given experimental condition, they are not linearly related to each other in terms of scale or between protocols; therefore, it is not valid to conclude that a reaction with twice the *max*/*K*_*D*_ of another reaction is “twice as bad”. For example, a comparison of two optimized probe-hydrolysis-based

qPCR diagnostic standards showed they have different inherent *max*/*K*_*D*_ ratios (**Supplementary Figure S1**).

### Identifying reaction outliers

Serial 10-fold dilutions of *E. coli* genomic DNA were used to establish a standard curve for a thermal-bias qPCR that targeted a section of the 16S rRNA gene. A replicate template sample of one of the dilutions was added to qPCR reagents that had been mis-mixed during setup (an ‘accidental’ second addition of the primer mix). Inspection of these reaction curves revealed that the mis-mixed samples exhibited delayed, yet productive, reactions compared to the correctly-mixed standards that contained the same amount of template (**Fig. 4A**).

**Figure 4.**
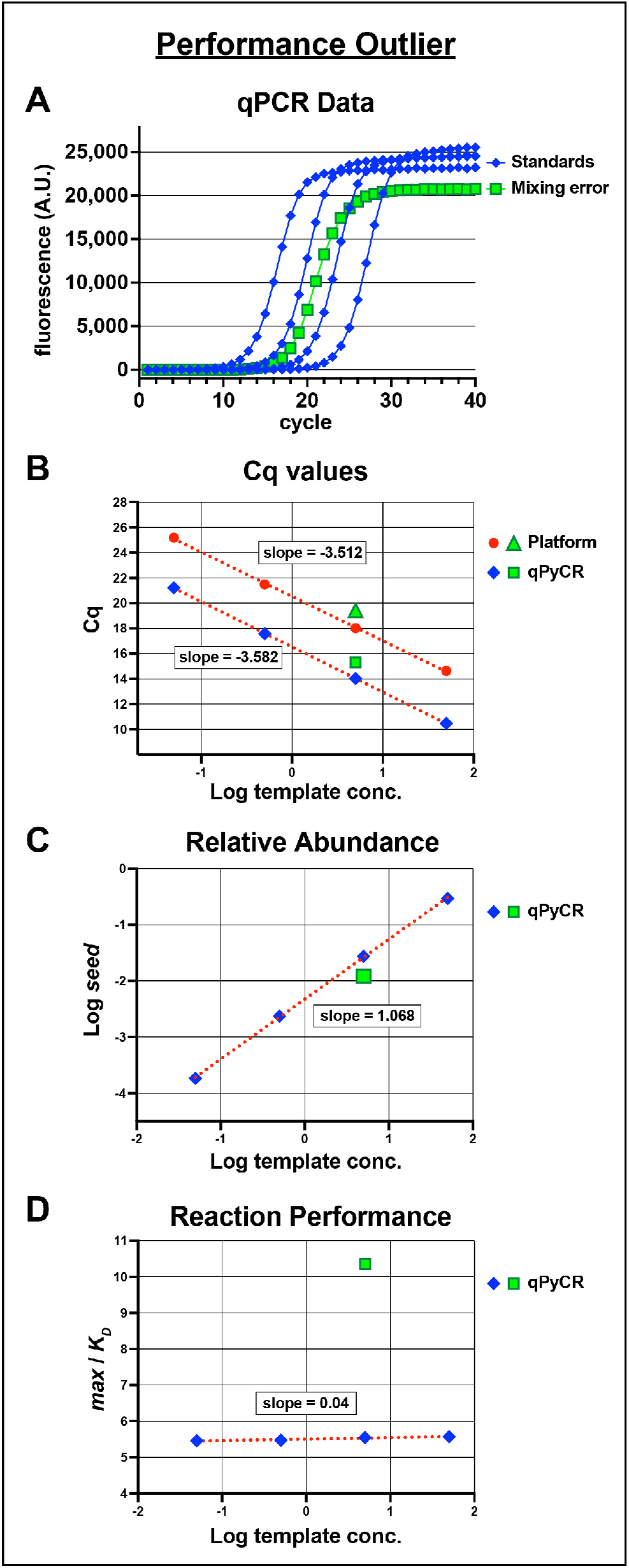
Identifying a performance outlier. A serial 10-fold dilution of genomic DNA template (50 to 0.05 μg/mL) was used for thermal-bias qPCRs. A separate qPCR reagent mixture was incorrectly prepared and used for an additional measurement of the 5 μg/mL standard. **A)** Reaction curves of the standards (blue diamonds) and the incorrectly-mixed sample (green squares), one of each of three replicates shown for clarity. The mis-mixed sample exhibited a delayed reaction, but generated abundant amplicons. **B)** Linear regression plots of Cq *vs*. Log(template concentration) of the standards and with the Cq of the mis-mixed sample also plotted (platform data: slope = -3.512, R^2^ = 0.9996; qPyCR data: -3.582, R^2^ = 0.9999), which correspond to per-cycle reaction efficiencies of 92.6% and 90.2%, respectively. **C)** Linear regression of *seed vs*. Log(template concentration) indicates a strong correlation (slope = 1.068, R^2^ = 0.9997). **D)** Template concentration-dependent reaction performance of the standards evaluated as *max*/*K*_*D*_ *vs*. Log(template concentration). The performance of these standards was notably consistent at ∼5.5, with a weak positive correlation (slope = 0.041, R^2^ = 0.931, Sy.x = 0.018). The mis-mixed sample performance values averaged ∼10.4 with a standard deviation of 0.18, which is extraordinarily deviant from the performance expectation derived from the standards (∼50-fold larger than the 95% prediction interval).

Consistent with delayed amplification, the averaged Cq and *seed* values indicated there was apparently less template in the mis-mixed reactions (**Fig. 4B** and **4C**). In contrast to the concentration-dependent drift in the *max*/*K*_*D*_ values of the standards shown in Fig. 3F, this thermal-bias protocol exhibited very similar performance at each template concentration (**Fig. 4D**). However, the performances of the mis-mixed samples were substantially out of range (**Fig. 4D**). When qPCR protocols are altered, the resulting *max*/*K*_*D*_ values can be assigned as new performance metrics for each condition (**Fig. S2**). Taken together, the qPyCR outputs provide a means to interrogate reaction performance relative to standards and they enable discrimination between faulty amplification and true changes in template abundance.

## Discussion and Conclusion

The qPyCR program provides a convenient tool to evaluate qPCR reaction quality. There are numerous assays that use Cq values for clinical diagnostics or for measuring template abundance (Untergasser et al., 2023); however, even with strict production quality controls, there is currently no way to distinguish apparent changes in target template abundance from inadvertent alterations in reaction quality. Different qPCR protocols yield different *max*/*K*_*D*_ ratios because of differences in per-cycle efficiencies, amplicon/probe signal strengths, and because reaction senescence mechanisms vary (Jansson and Hedman, 2019). With these considerations in mind, it is envisioned that qPCR reaction curve shape analysis can be integrated into production validation pipelines, to increase measurement confidence, or to evaluate the performance of alternative measurement protocols (Gaby and Buckley, 2017; Gohl et al., 2016; Kahsen et al., 2024; Kolm et al., 2020; Pearson et al., 2022).

Although Cq analysis is well-vetted for its quantification ability, it relies on two underlying assumptions. The first is that the fastest amplification regions yielding near-exponential data should log-transform into strictly linear data (Ramakers et al., 2003); however, per-cycle PCR efficiency is not necessarily constant (Carr and Moore, 2012; Hampton et al., 2025). The second assumption is that the crossing point of a threshold with a log-linear regression curve represents a fluorescence signal at that point (Higuchi et al., 1993; Pfaffl, 2001), but qPCR data are histograms of measurements at the ends of each cycle and there is not a linear relationship between their log-transforms. While these caveats do not substantially impact the utility of Cq-based analyses (Bustin et al., 2025; Ruiz-Villalba et al., 2021), they can reinforce misconceptions of the underlying amplification process. For these reasons, alternative models for qPCR data analysis are useful for interrogating reaction performance (Patrone et al., 2020).

Template-concentration-dependent changes in reaction performance were observed for a commercial quantification standard (**Fig. 3F**). Similar changes have been observed by us and other groups for unrelated dilution-series experiments (Ruijter et al., 2019). Template-dependent changes in reaction performance cause the resulting Cq *vs*. log-template plots to change their slopes and therefore the calculated per-cycle efficiencies. The co-amplification of ‘artifact’ products contribute to this phenomenon and they can be monitored and accounted for when accurate quantification is desired (Ruijter et al., 2019). Another contribution to template-concentration-dependent changes in reaction performance arises from repeated priming on the initial target (Green et al., 2015). Such repeated targeting can be initiated either by the primers themselves or by previously generated amplicon strands (which also prevents those strands from being primed). Evidence supporting this behavior lies in the nearly concentration-independent reaction performance observed using thermal-bias PCR (**Fig. 4D**), which avoids retargeting so that the observed amplification signals are primarily from chain-reactions. Thus, the fluorescence data can contain additional artifactual exponential and linear signals that alter the apparent efficiency and reaction profile shapes.

qPyCR is presented in a cell-based format that allows each operational stage to be monitored. This format is intended to encourage the development and implementation of alternative approaches to baseline adjustment, Cq calling, or data fitting. Most commercial qPCR platforms have project setup dialogs that assign standards and allow for the separation of different targets or reaction types, so an integration of a reaction performance metric for a given condition should be fairly straightforward to implement. It is anticipated that qPyCR will make global fitting more approachable to casual users of qPCR and facilitate diagnostics and protocol optimization.

## Supporting information

Primer details

## Code Availability

The modular Jupyter .ipynb notebook (https://jupyter.org/) and a stand-alone Python .py script are available in a public GitHub repository (https://github.com/sdmoore-labs/qPyCR). The GitHub README.md file contains a link to run the program in a free online Juptyer Binder without local installation. A test_data.csv file and a Test_Data_RDES_amplification.tsv file are located in the GitHub qPyCR ‘examples’ directory. Code archives are present on the Zenodo server (https://doi.org/10.5281/zenodo.21115342).

## Author contributions

Sean Moore is the sole contributor.

## Acknowledgements

The raw and platform-adjusted qPCR data used in Figs. 2E-G, 3D, and S1 were kindly provided by Veronica Urdaneta and Taj Azarian (UCF Burnett School of Biomedical Sciences, Orlando, FL). The name qPyCR was coined by Andrew Kettring.

