## Supplementary Data SD1 for "qPyCR: A qPCR Data Analyzer for Template Quantification and the Detection of Performance Outliers": ReadMe for NEBNext Standards Data.rtf

NEBNext Standards Data
NEBNext_Standards_All.csv = Raw fluorescence data from three replicate measurements for each sample.
NEBNext_One_Of_Each.csv = One selection from each replicate.

target concentrations:
A1-A3 = 100 pM
B1-B3 = 10 pM
C1-C3 = 1 pM
D1-D3 = 0.1 pM
E1-E3 = 0.01 pM
F1-F3 = 0.001 pM

Reagents:
NEBNext® Library Quant Kit for Illumina NEB #E7630S/L according to the manufacturer's instructions. Kit details and data reference:
https://www.neb.com/en-us/products/e7630-nebnext-library-quant-kit-for-illumina

Platform:
Bio-Rad CFX Opus
CFX Maestro Version 5.3.022.1030.
