## Supplementary Data SD1 for "qPyCR: A qPCR Data Analyzer for Template Quantification and the Detection of Performance Outliers": ReadMe for Performance Outlier.rtf

Performance Outlier Data
Performance_Outlier_All.csv = Raw fluorescence data from three replicate measurements for each sample.
Performance_Outlier_One_Of_Each.csv = One selection from each replicate.

Samples:
50_1 - 50_3		= 50 ìg/mL
5_1 - 5_3		= 5 ìg/mL
5_1_me - 5_3_me	= 5 ìg/mL in mis-mixed reagent
0.5_1 - 0.5_3		= 0.5 ìg/mL
0.05_1 - 0.05_3	= 0.05 ìg/mL

Reagents and procedure
E. coli genomic DNA diluted in carrier buffer (0.1 mg/mL yeast tRNA, 0.05% Tween-20, 10 mM Tris-Cl, 0.1 mM EDTA, pH 8.0).

1X SsoFast™ EvaGreen® Supermix (cat. 1725201), 0.5 ìM each primer, and 1/20th volume of 50-0.05 ìg/mL E. coli genomic DNA (2,500 - 2.5 pg/ìL in final mix), 20 ìL of mix in each well.

The mis-mixed sample contained an additional delivery of primers to a master mix, so the resulting final mixture contained ~0.7 X SsoFast Supermix, and ~0.7 ìM each primer.

Thermal cycling:
98 °C	2 min
2X:
98 °C	10 sec
50 °C	5 min
40X:
98 °C	10 sec
78 °C	1 min
read FL

Platform:
Bio-Rad CFX Real-time System
CFX Manger version 3.1.1517.0823
