## Supplementary Data SD1 for "qPyCR: A qPCR Data Analyzer for Template Quantification and the Detection of Performance Outliers": ReadMe for Primer performance Data.rtf

Primer Performance Data
Primer_Performance_All.csv = Raw fluorescence data from three replicate measurements for each sample.
Primer_Performance_One_Of_Each.csv = One selection from each replicate.

Primer pairs:
B3-B5 = Pair_1
C3-C5 = Pair_2
D3-D5 = Pair_3

Reagents and procedure
E. coli genomic DNA diluted in carrier buffer (0.1 mg/mL yeast tRNA, 0.05% Tween-20, 10 mM Tris-Cl, 0.1 mM EDTA, pH 8.0).

1X SsoFast™ EvaGreen® Supermix (cat. 1725201), 0.5 ìM each primer, 5 ìg/mL E. coli genomic DNA (0.25 ìg/mL in mix), 20 ìL of mix in each well.

Thermal cycling:
98 °C	2 min
2X:
98 °C	10 sec
50 °C	5 min
40X:
98 °C	10 sec
78 °C	1 min
read FL

Platform:
Bio-Rad CFX Real-time System
CFX Manger version 3.1.1517.0823
